## Supplementary Material for "Genetic impacts on within-pair DNA methylation variance in monozygotic twins capture gene-environment interactions and cell-type effects"

### 1. Supplementary methods and results

#### 1.1 Methods comparison with simulated data

To compare the method performance of Twins1 to Twins6 methods. We transformed simulated DNA methylation beta value to DNA methylation M value (M=log2(beta/1-beta)). We repeated the simulations 1000 times and calculated how frequently each method could detect the significant genotype effects on the intra-twin differences with the P value of genotype term surpass P<0.05. We observed that the Twins5 and Twins6 methods show high false positive rates when the errors follow non-normal distributions, and Twins3 and Twins4 methods can barely detect any vmeQTL effects. Twins1 method shows the highest discovery rate when the genetic effects of GxE interactions are lower than 0.2 across all data distributions. However, The FPR of method Twins1 was greater than 0.05 when the errors followed non-normal distributions (Supplementary Figure 2A) and its discovery rate was lower than that obtained using beta values (Supplementary Figure 1B-C and 2B-C)

#### 1.2 Suprious vmeQTLs and meQTLs exclusion

The majority of tested CpG sites exhibit correlation between the mean intra-twin-pair methylation level and difference greater than 0.3 (Spearman’s rank, Supplementary Figure 3B), therefore, the meQTLs or the SNPs in linkage disequilibrun (LD) with meQTLs could be spurious vmeQTLs, and vice versa. Therefore, for each CpGs that were affected by vmeQTLs and meQTLs (2,230 CpGs for cis and 460 CpGs for trans effects), we carried out the LD clumping with Plink (window = 500kb, r2 = 0.2) of all SNPs that were identified as vmeQTLs and meQTLs according to the P-values. Specifically, the CpGs affected by 1/ cis-vmeQTLs and cis-meQTLs, 2/ cis-vmeQTLs and trans-meQTLs, 3/ trans-vmeQTLs and cis-meQTLs, and 4/ trans-meQTLs and trans-meQTLs. Following LD clumping, if vmeQTL SNPs fell in LD clumps together with meQTLs, then these vmeQTLs were removed and vise versa. In these steps, we observed 914 vCpGs for cis and 457 vCpGs for trans. Altogether with the CpGs affected by vmeQTLs only (107 cis-vCpGs and 199 trans-vCpGs), 1,021 cis-vCpGs and 656 trans-vCpGs involved in 1,703 associations were kept for further filtering. For 1,703 associations, we evaluated whether the vmeQTL SNP is still associated with intra-twin methylation difference by adding vmeQTL SNP as a covariate, 5 vmeQTL-vCpG associations were removed. No associations were excluded in the later sensitivity analysis. In total, we have observed 1,698 vmeQTL-vCpG associations (1,017 vCpGs for cis and 656 vCpGs for trans) for further exploration.

#### 1.3 The enrichment of transcript factor binding sites of 1,698 associations in TwinsUK.

In terms of the enrichment analysis of each transcript factor binding sites of the 1,698 associations in TwinsUK, we observed that *trans* vCpGs are enriched in SUZ12 (OR = 2.00, CI [1.46, 2.68], P=1.73e-5) and EZH2 (OR = 1.56, CI [1.06, 2.23], P=1.86e-2), which are the core subunits of polycomb repressive complex 2 (PRC2) (Supplementary Figure 6). PRC2 is evolutionarily conserved and plays an important role in cellular proliferation and histone methyltransferase activity. This finding is consistent with previous studies that propose that *trans*-regulatory effects play an important role in intraspecies expression variance in natural selection and evolution.

### 2. Supplementary figures

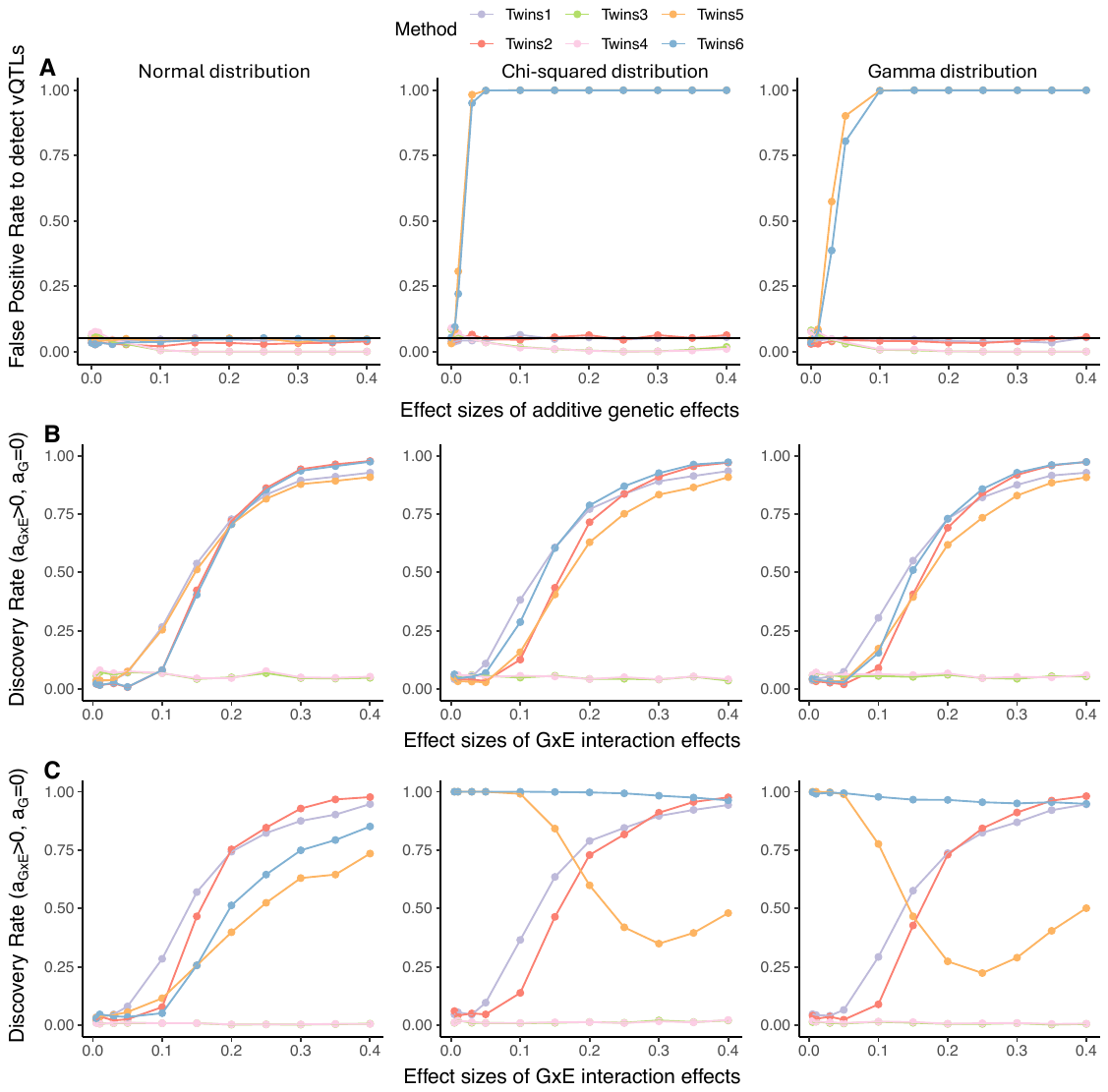

**Supplementary Figure 1. Method performance with simulated DNA methylation beta value.** A) False positive rates estimation when there are no GxE interaction effects (a_G_>=0, a_GxE_=0). Discovery rates estimation when there are GxE interaction effects, including two parameter settings B) a_G_=0, a_GxE_>0, and C) a_G_=0.1, a_GxE_>0. The three columns represent the errors follow Normal (left), Chi-squared (middle), and Gamma distributions (right). The six models (Twins1-6) included Twins1: res(abs) ~ genotype; Twins2: res(abs)2 ~ genotype; Twins3: res(abs)/res(mean) ~ genotype; Twins4: res2(abs)/res2(mean) ~ genotype; Twins5: res(abs) ~ genotype + res(mean); Twins6: res2(abs) ~ genotype + res(mean).

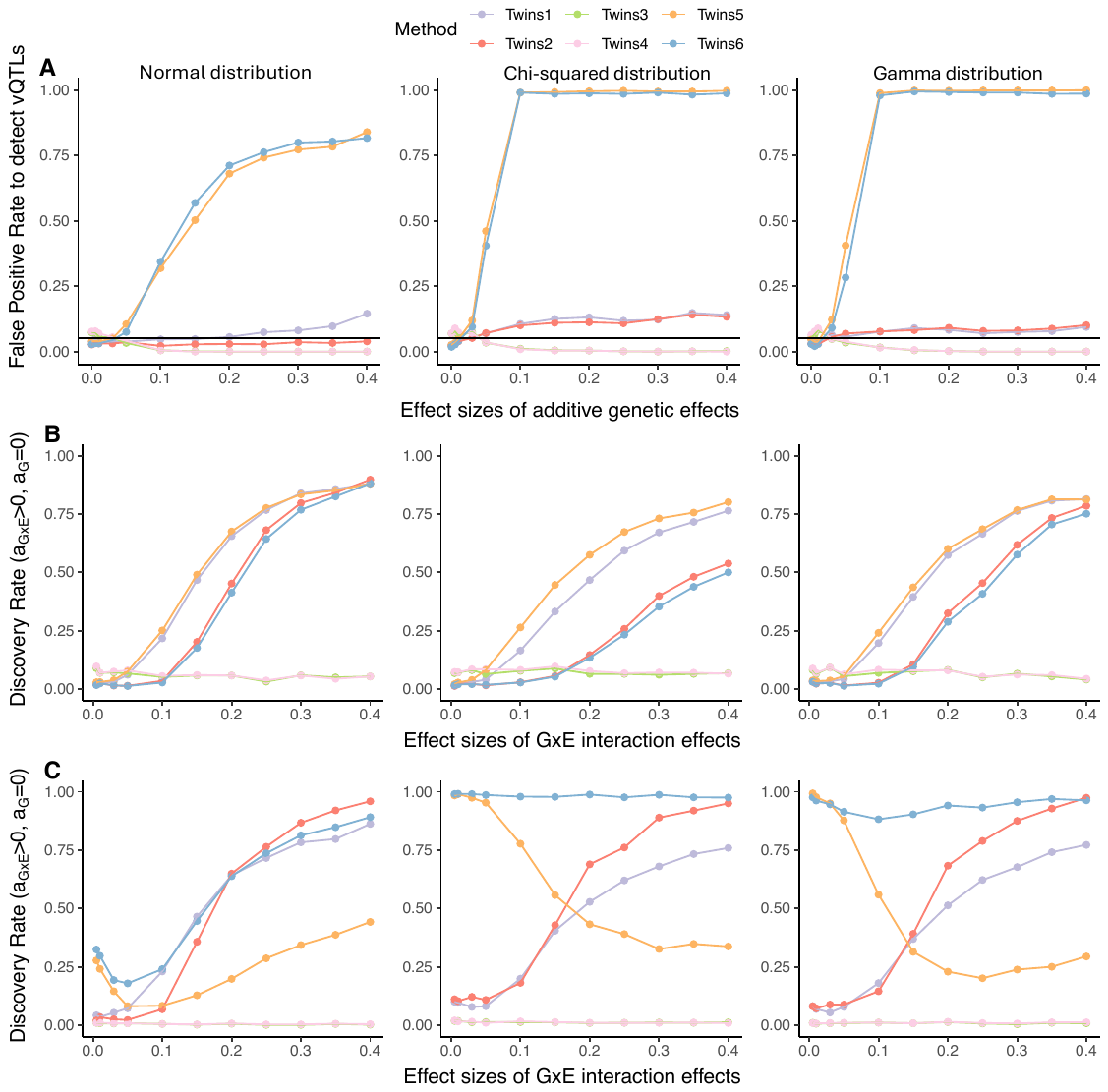

**Supplementary Figure 2. Method performance with simulated DNA methylation M value.** A) False positive rates estimation when there are no GxE interaction effects (a_G_>=0, a_GxE_=0). Discovery rates estimation when there are GxE interaction effects, including two parameter settings B) a_G_=0, a_GxE_>0, and C) a_G_=0.1, a_GxE_>0. The three columns represent the errors follow Normal (left), Chi-squared (middle), and Gamma distributions (right). The six models (Twins1-6) included Twins1: res(abs) ~ genotype; Twins2: res(abs)2 ~ genotype; Twins3: res(abs)/res(mean) ~ genotype; Twins4: res2(abs)/res2(mean) ~ genotype; Twins5: res(abs) ~ genotype + res(mean); Twins6: res2(abs) ~ genotype + res(mean).

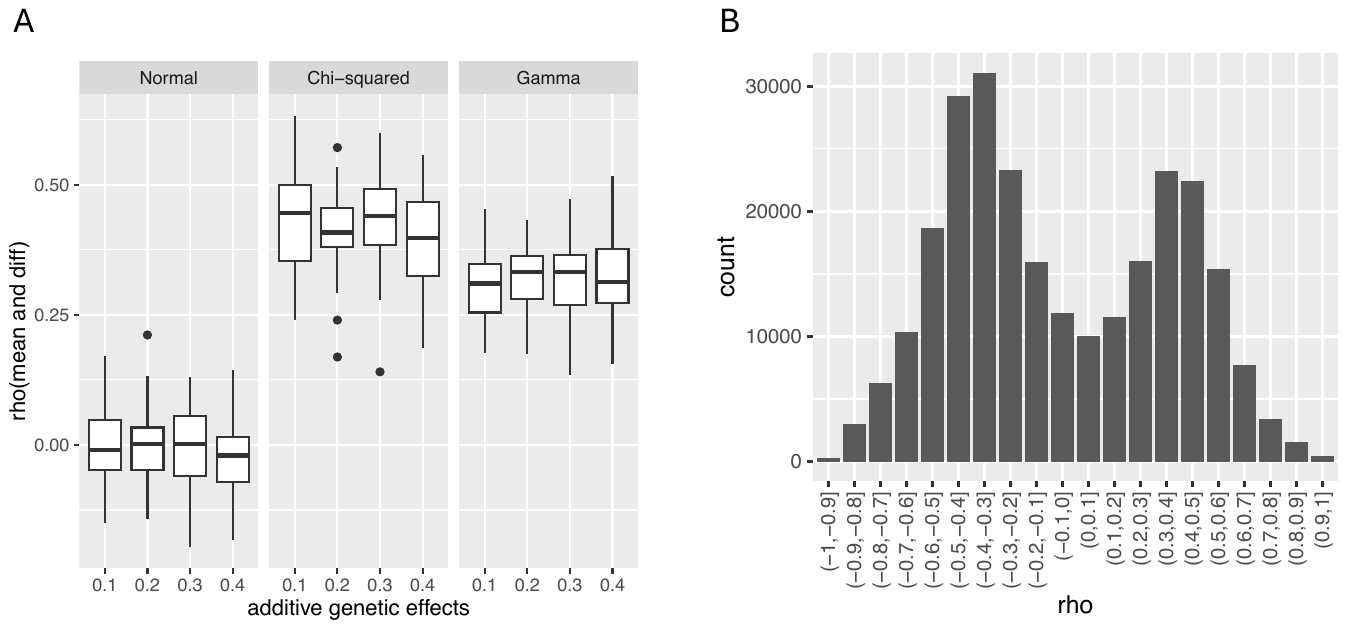

**Supplementary Figure 3. The correlation between the mean within-twin pair phenotype level and intra-twin pair phenotype difference, estimated within each twin pair.** A) The correlation of 1000 simulated CpGs under the scenario where there is no GxE interaction on DNA methylation. Three columns represent the errors follow Normal (left), Chi-squared (middle), and Gamma (right) distributions, respectively B) The correlations of all tested CpGs (~261K) across 355 pairs of twins.

**Supplementary Figure 4. The vmeQTLs detection workflow in TwinsUK.**

*****N represents the number of vCpGs in cis or trans. 2230 cis-vCpGs include the CpGs affected by cis-vmeQTLs and meQTLs (including cis- and trans-meQTLs), and 460 trans-vCpGs represents CpGs affected by trans-vmeQTLs and meQTLs (including cis- or trans-meQTLs).

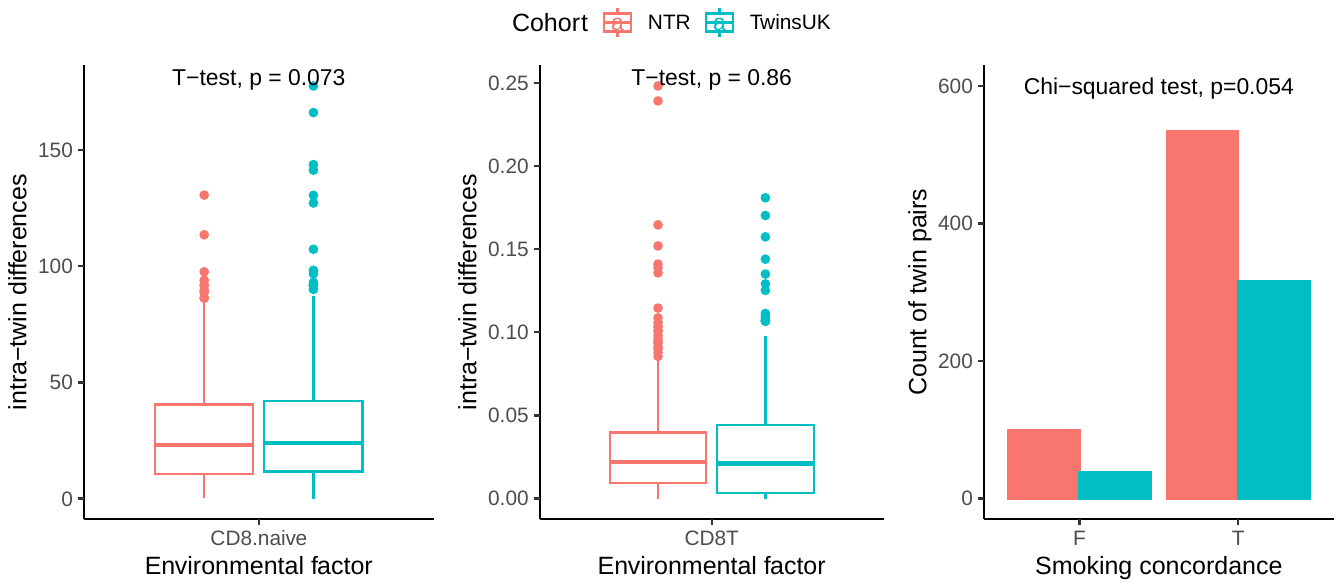

**Supplementary Figure 5. The intra-twin pair differences of naïve CD8+ T cells, CD8+ T cells, and the consistency of smoking status in TwinsUK and NTR.**

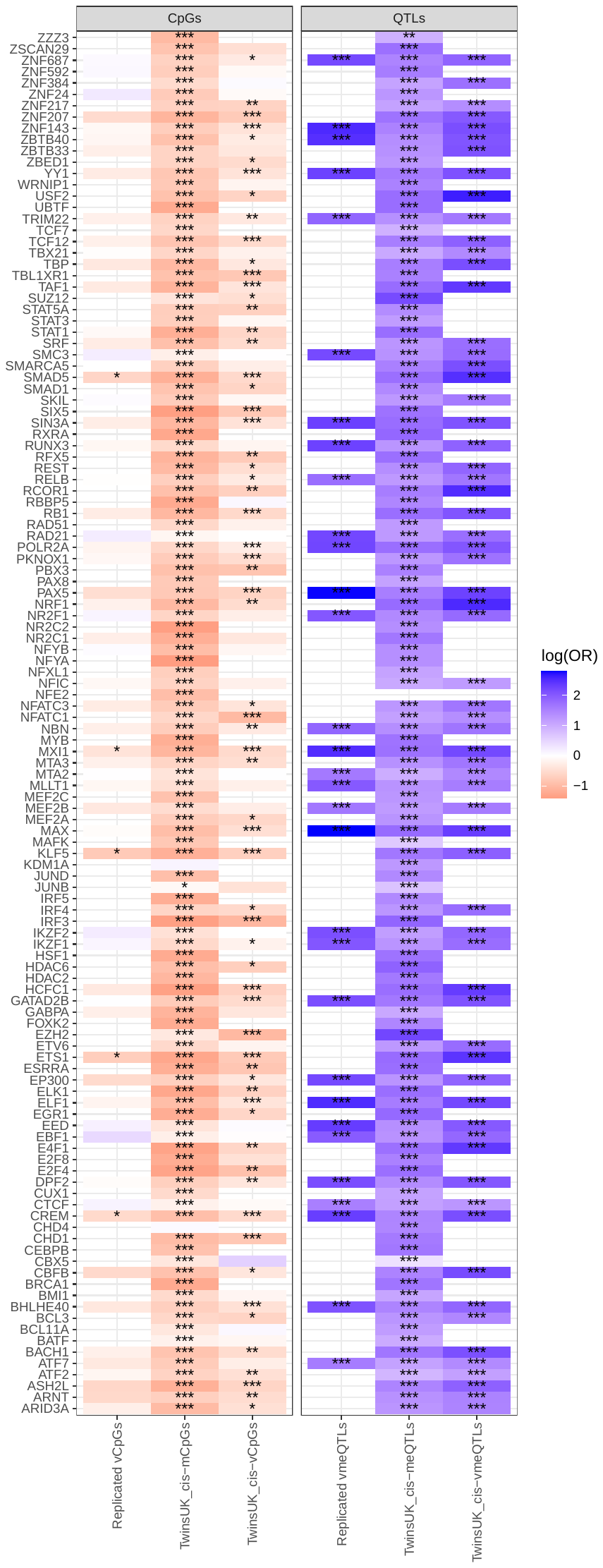

**Supplementary Figure 6. The genomic distributions of CpGs and QTLs for each TFBS.** The omitted or blanked regions have fewer than 10 signals. The number of stars represents the FDR of the enrichment or deletion, *** FDR<0.001, ** FDR<0.01, and * FDR<0.05.

### 3. Cohort-specific acknowledgements

#### 3.1 TwinsUK

TwinsUK is funded by the Medical Research Council (MRC), Wellcome LEAP, Wellcome Trust, EPSRC, BBSRC, Versus Arthritis, European Commission, Chronic Disease Research Foundation (CDRF), Zoe Ltd, the National Institute for Health and Care Research (NIHR) Clinical Research Network (CRN) and Biomedical Research Centre based at Guy’s and St Thomas’ NHS Foundation Trust in partnership with King’s College London.

#### 3.2 Netherland Twin Registry

Netherlands Twin Registry acknowledge funding from the Netherlands Organization for Scientific Research (NWO): Biobanking and Biomolecular Research Infrastructure (BBMRI-NL, NWO 184.033.111) and the BBRMI-NL-financed BIOS Consortium (NWO 184.021.007), NWO Large Scale infrastructures X-Omics (184.034.019), Genotype/phenotype database for behaviour genetic and genetic epidemiological studies (ZonMw Middelgroot 911-09-032); Netherlands Twin Registry Repository: researching the interplay between genome and environment (NWO-Groot 480-15-001/674); the Avera Institute, Sioux Falls (USA), and the National Institutes of Health (NIH R01 HD042157-01A1, MH081802, Grand Opportunity grants 1RC2 MH089951 and 1RC2 MH089995); epigenetic data were generated at the Human Genomics Facility (HuGe-F) at ErasmusMC Rotterdam. DIB acknowledges the Royal Netherlands Academy of Science Professor Award (PAH/6635).

#### 3.3 The MRC National Survey of Health and Development or1946 British birth cohort

The U.K. Medical Research Council provides core funding for the MRC National Survey of Health and Development (MCUU00019/1). KKO is supported by the Medical Research Council (MCUU00006/2). We acknowledge study members for their lifelong participationand past and present members of the study teams, including members of the MRCEpidemiology unit in Cambridge, who helped to collect and process the data.
